# Alzheimer’s disease Phospholipase C-gamma-2 (PLCG2) protective variant is a functional hypermorph

**DOI:** 10.1101/409706

**Authors:** Lorenza Magno, Christian B Lessard, Marta Martins, Pedro Cruz, Matilda Katan, Jamie Bilsland, Paramita Chakrabaty, Todd E Golde, Paul J Whiting

**Affiliations:** UCL Alzheimer’s Research UK, Drug Discovery Institute, London, UK; Department of Neuroscience, Center for Translational Research in Neurodegenerative Disease, and McKnight Brain Institute, College of Medicine, University of Florida, Gainesville, FL, United States; Research Department of Structural and Molecular Biology, University College London, London, UK; present address Instituto de Medicina Molecular - João Lobo Antunes, Faculdade de Medicina de Lisboa, Lisbon, Portugal; Dementia Research Institute, UCL, London, UK

## Abstract

Recent Genome Wide Association Studies (GWAS) have identified novel rare coding variants in immune genes associated with late onset AD (LOAD). Amongst these, a polymorphism in Phospholipase C-gamma 2 (PLCG2) P522R, has been reported to be protective against LOAD. PLC enzymes are key elements in signal transmission networks and are potentially druggable targets. PLCG2 is highly expressed in the hematopoietic system. Hypermorphic mutations in PLCG2 in humans have been reported to cause autoinflammation and immune disorders, suggesting a key role for this enzyme in the regulation of immune cell function.

We confirmed that PLCG2 expression is restricted primarily to microglia in both the healthy and AD brain. Functional analysis of the P522R variant in heterologous systems demonstrated a small hypermorphic effect of the mutation on enzyme function. PLCγ2 is therefore a potential target for modulating microglia function in AD, and a small molecule drug that weakly activates PLCγ2 may be one potential therapeutic approach.

**SUMMARY:** The *PLCG2* P522R variant is protective against Alzheimer’s disease (AD). We show that PLCG2 is expressed in CNS-resident myeloid cells, and the P522R polymorphism weakly activates enzyme function. These data suggest that activation of PLCG2 and not inhibition could be therapeutically beneficial in AD.

## INTRODUCTION

Alzheimer’s disease is the most common neurodegenerative disorder, and the leading cause of dementia. Late onset AD (LOAD) is genetically complex, and known susceptibility loci only explain a proportion of disease heritability (Gatz et al., 2006). Large scale genetic studies have led to the identification of several new susceptibility genes associated with LOAD. These genetic data, together with analysis of biological pathways, implicate processes related to immune response in the aetiology of LOAD, and point to the immune system as a prime target for therapeutic approaches (Zhang et al., 2013; Jones et al., 2015; Efthymiou and Goate, 2017). Amongst the newly discovered polymorphisms, rare variants in microglial-related genes including Triggering Receptor Expression on Myeloid cell-2 (TREM2), ABI family member 3 (ABI3) and Phospholipase C-gamma-2 (PLCG2) have been described (Guerreiro et al., 2013; Sims et al., 2017). Notably, the PLCG2 missense variant Pro522Arg (P = 5.38 × 10^−10^, OR = 0.68) was associated with decreased risk of LOAD (Sims et al., 2017).

PLCG2 belongs to the family of phospholipase C-gamma and encodes an enzyme (PLCγ2) that cleaves the membrane phospholipid PIP2 (1-phosphatidyl-1d-myoinositol 4,5-bisphosphate) to secondary messengers IP3 (myoinositol 1,4,5-trisphosphate) and DAG (diacyl-glycerol) which further propagate a wide range of downstream signals. PLCγ2 shares high structural and mechanistic overlap with the other member of the PLCγ family, PLCγ1 (Koss et al., 2014). Although both enzymes are important for regulation of specific responses of specialised cells of the immune system, they show different cell-type expression and are relevant in very different medical conditions (Koss et al., 2014). PLCG1 is ubiquitously expressed, and mutations are associated with some forms of cancers, such as cutaneous T cell lymphoma (Bunney and Katan, 2010; Vaqué et al., 2014), but also neuropsychiatric disorders (Yang et al., 2017). PLCγ2 is predominantly expressed in the bone marrow and lymphoid organs (Human Protein Atlas available from www.proteinatlas.org; Mao et al., 2006), and PLCG2 variants cause inherited immune disorders designated as PLAID (PLCG2-associated antibody deficiency and immune dysregulation; Ombrello et al., 2012) and APLAID (autoinflammatory PLAID; Zhou et al., 2012).

In immune cells, PLCγ2 has been implicated in signalling pathways downstream of B and T cell receptors, and it is thought to modulate the functions of macrophages, platelets, mast cells, neutrophils and NK cells through Fc receptor signalling (Wang et al., 2000). PLC enzymes belong to the same interaction network as Triggering Receptor Expression on Myeloid cell-2 (TREM-2; Sims et al., 2017) and may be directly involved in its signalling pathway (Ford and McVicar, 2009; Xing et al., 2015). TREM2 is a transmembrane receptor expressed on the membrane of myeloid cells, and an important susceptibility gene for AD (Guerreiro et al., 2013). In osteoclasts, activation downstream Trem2 initiates a cascade of events, including phosphorylation and activation of PLCγ2, amongst other proteins (Peng et al., 2010). In microglia, these pathways promote several cellular responses including survival, proliferation, phagocytosis, enhanced secretion of cytokines and chemokines (Colonna and Wang, 2016) and are potentially involved in neurodegenerative processes.

Genomic studies can accelerate the pathways to novel therapeutics by suggesting promising translational targets, and informing the type of approach to be undertaken. The identification of naturally occurring genetic variants with protective effects against diseases represents a valuable potential resource for drug development (Harper et al., 2015; Butler et al., 2017). The P522R polymorphism lies within the regulatory domain of PLCγ2, but its effect on enzyme function is unknown. Given that the variant is protective against LOAD, understanding its effect is fundamental in determining whether a small molecule inhibitor or activator/stabilizer is required as a therapeutic.

PLCγ2 spatial expression in resident brain immune cells and in AD is unexplored. Therefore, we first characterised its expression in the brain. We found that *Plcg2* mRNA mainly co-localises with microglia-specific markers in healthy brain tissue, as well as in microglia near amyloid plaques in an APP mouse model. Additionally, functional characterisation of the AD protective variant PLCγ2 P522R revealed a small increase in activity compared to wild type enzyme. PLCγ2 is therefore a potential target for modulating microglia function in AD, and a small molecule drug that activates PLCγ2 may be one potential therapeutic approach.

## RESULTS AND DISCUSSION

### PLCG2 is expressed in microglia cells throughout the brain

Recent transcriptomic datasets suggest prominent *Plcg2* expression in the CNS resident myeloid cells of the mouse cortex (Bennett et al., 2016; Tasic et al., 2016.; Zeisel et al., 2015), however spatial mapping of the protein and mRNA in brain tissue has not been described. To better understand the involvement of this protein in AD, we assessed PLCγ2 expression in human cortical tissue. We detected PLCγ2 immunoreactivity in putative microglia of both white and grey matter, in the human frontal cortex (Figure 1A). Due to technical difficulties with the antibodies used in this study (in one case the antibody is no longer commercially available; in other cases, antibodies gave unspecific or no signal upon IHC on fixed human post-mortem brain or mouse tissue) we turned to *in situ* hybridization (RNAScope, Wang et al., 2012) to further investigate the expression patterns of *Plcg2* across several regions of the mammalian brain. Since PLCγ1 seems to be expressed ubiquitously in mammalian tissues, while PLCγ2 is thought to be restricted to the hematopoietic and immune system (Koss et al., 2014), we sought to determine the distribution of these two PLCs in the CNS. We found that *Plcg1* and *Plcg2* mRNAs show complementary expression patterns, with very little degree of overlap in the adult mouse brain (Figure 1B). Labelling for *Plcg1* was detected in several brain cells, including large nuclei of putative neurons of the cortex (Figure 1B). Conversely, *Plcg2* reaction product was found in sparse cells, in cortical as well as subcortical regions, including the olfactory bulb, neocortex, hippocampus CA1, and substantia nigra (Supplementary Figure 1B).

**Figure 1.**
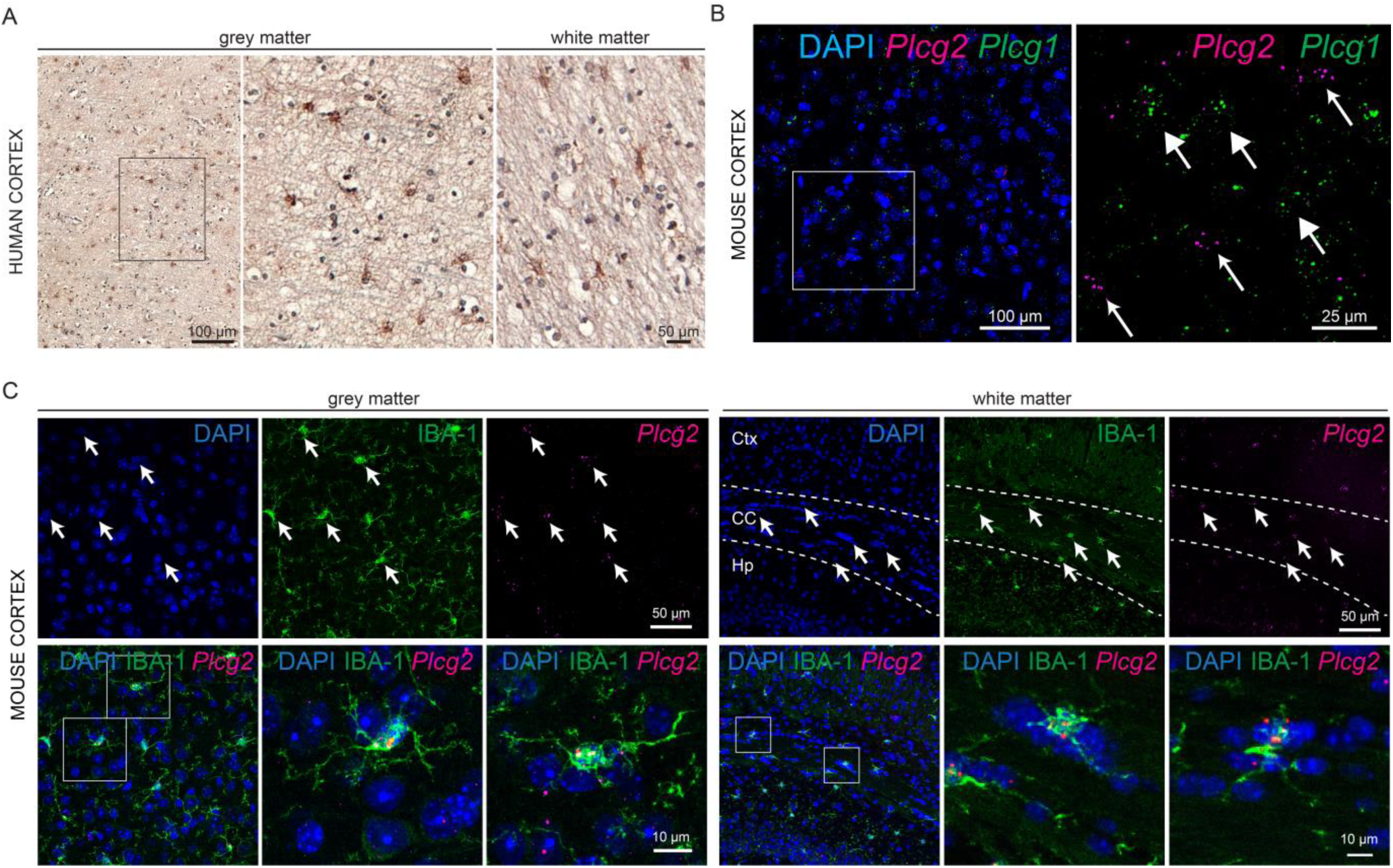
PLCγ2 expression in the human and mouse cortical microglia A) IHC for PLCγ2 on the grey and white matter of the human prefrontal cortex. B) Multiplex RNAScope for *Plcg2* and *Plcg1* on the adult mouse cortex. Small and big arrows point to RNASCope reaction for *Plcg2* and *Plcg1* respectively. C) RNAScope for *Plcg2* followed by IHC for IBA-1 on the grey and white matter of the adult mouse cortex. Arrows point to co-expression of the two markers. Ctx, cortex; CC, corpus callosum; Hp, hippocampus.

In line with the single-cell RNA-Seq profiling databases of the mouse cortex (see above), we confirmed a high degree of co-localisation between *Plcg2* mRNA and the microglia-specific marker IBA-1 in cortical grey and white matter of the adult mouse brain (Figure 1A), and we further validated co-expression between the two markers in subcortical brain regions, including the thalamus and the cerebellum (data not shown).

We further assessed co-localisation between *Plcg2* and other glia/endothelial cell markers, and found only occasional co-expression with markers for astrocytes (*Glast*, Supplementary Figure 2A), endothelial cells (*Pecam-1*, Supplementary Figure 2B), and oligodendrocyte-lineage cells (*Olig2*, Supplementary Figure 2C).

In addition to a generally scattered labelling in most brain areas, we found intense *Plcg2* RNAScope reaction product in the granule cell layer of the hippocampal dentate gyrus, where the *Plcg2* signal co-localises with the neuronal marker NeuN (Supplementary Figure 2D). The role of Plcg2 in these neurons is unknown. By analogy with Plcg1 function in neurons, expression of Plcg2 in granule cells may be relevant for synaptic transmission and plasticity via induction of hippocampal long-term potentiation LTP (Yang et al., 2017; Horn et al., 2013). Given the involvement of the dentate gyrus in pattern separation - distinction of closely related memories-Plcg2 expression might be important for mnemonic functions. Interestingly, to date no deficits in learning and memory have been described in Plcg2 -/- mice. Rare co-labelling with NeuN was also detected in other subcortical regions (not shown). In summary, Plcg2 expression is primarily restricted to microglia cells across the mouse brain, with the exception of the expression seen in granule cells in the dentate gyrus.

### PLCG2 in microglia in Alzheimer’s disease

PLCG2 RNA upregulation has been reported in cortical tissue of LOAD patients (FC, 1.35; p = 0.0028, Allen et al., 2016) and in transgenic mice carrying mutations associated with early AD (APP KM670/671 NL, PNSEN1 M146V) or overexpressing the human tau-4R/2N isoform (P301L) (Castillo et al., 2017; Matarin et al., 2015). However, normalisation to microglia-specific genes (eg. Abi3) and data from single-cell profiling of CNS immune cells (Sims et al., 2017; Keren-Shaul et al., 2017; Mathys et al., 2017), suggest that PLCG2 upregulation in bulk tissue is mostly related to microgliosis occurring as a consequence of neurodegeneration.

Recent findings point to microglia heterogeneity in neurodegeneration (Keren-Shaul et al., 2017; Krasemann et al., 2017). To determine whether heterogeneity in Plcg2 expression might be observed in microglia in AD, we carried out *in situ* hybridization for *Plcg2* on the brain of a transgenic mouse model overexpressing the human APP with the Swedish (KM670/671NL) and Indiana (V717F) mutations (TgAPPswe/ind, Chishti et al., 2001). *Plcg2* appeared to be equally expressed in microglia surrounding the plaques in the cortical regions of 6 months old transgenic mice (Figure 2A) as compared to expression in microglia in control non-transgenic littermates (Figure 2B).

**Figure 2.**
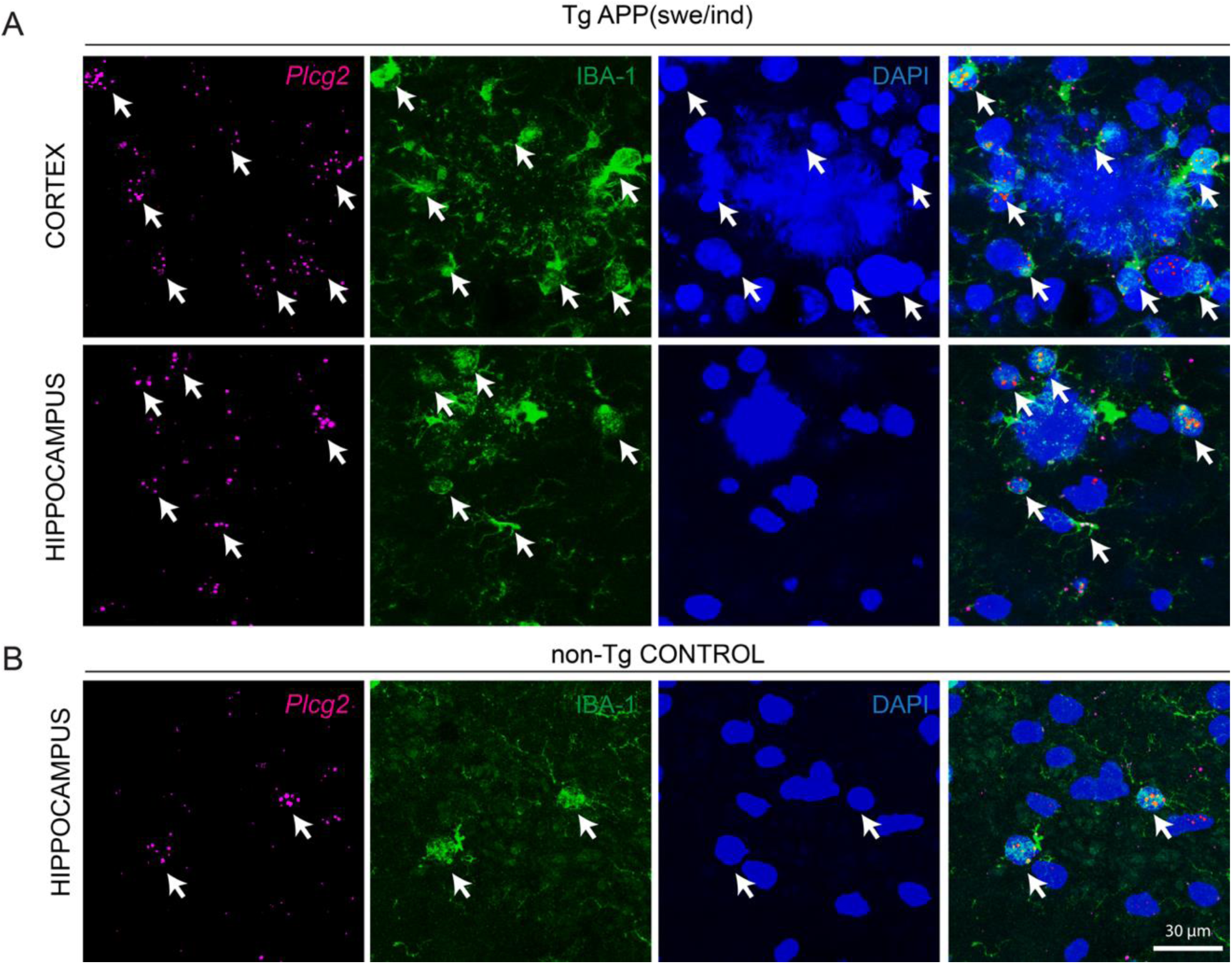
*Plcg2* mRNA localises in microglia at amyloid plaques in a mouse model of AD. A) Representative images of RNAScope labelling for *Plcg2* in adult mouse neocortex and hippocampus of *APP* transgenic mice. Note expression of *Plcg2* in IBA-1 labelled microglia surrounding the plaques (DAPI). Arrows point to co-expression of IBA-1 and *Plcg2*. B) Representative images of RNAScope labelling for *Plcg2* in adult mouse hippocampus of *a* non transgenic control mouse. Arrows point to co-expression of IBA-1 and *Plcg2*

Next, we assessed whether Plcg2 co-localise with the myeloid Trem2 receptor in the brain. In line with protein-protein interaction network models (Sims et al., 2017), we detected co-expression of *Plcg2* mRNA with *Trem2* mRNA in the cell soma and processes of putative cortical and hippocampal microglia (Supplementary Figure 3). mRNA localisation in the processes points to a possible requirement of local translation, for quick responses to environmental changes, and a mechanism to detect neural tissue damage. This pathway could be critical for microglia activity in neurodegenerative conditions.

### AD associated PLCG2 P522R variant shows a weak hypermorphic activity

Germline deletion and point mutations in PLCG2 cause complex immune disorders and autoinflammatory disease in humans (Ombrello et al., 2012; Zhou et al., 2012). Similarly, two Plcg2 mutations in mice (Ali5, D993G; Ali14, Y495C) lead to severe autoinflammation and antibody deficiencies (Yu et al., 2005; Abe et al., 2011; Everett et al., 2009). Activation of the PLC family of proteins catalyses the hydrolysis of phosphatidylinositol 4,5-bisphosphate producing inositol 1,4,5-trisphosphate (IP3) and diacyl-glycerol. Binding of IP3 on its receptor induces intracellular Ca2+ release from the Ca2+ stored in endoplasmic reticulum. The effects of the above mutations on enzyme function have been determined by quantifying the production of IP3 or the release of intracellular Ca2+ following stimulation with epidermal growth factor (EGF) in heterologous cell system *in vitro* (Koss et al, 2014).

To investigate the effect of the AD protective variant P522R on PLCγ2 enzyme function we firstly, measured ^3^[H]inositol phosphate production in a radioactive assay as described by Everett et al., (2009) upon transient transfection with pTriEx4-PLCG2 (common, “wild type” variant; PLCG2 522P) construct; the AD associated (P522R) construct; and with the D993G mutation (Ali5, Yu et al., 2005) construct in COS7 cells. PLC activity after stimulation with EGF showed a 1.24 ± 0.06 fold increase (mean value ± SD over 3 independent experiments, normalised to PLCγ2 WT activity) upon transfection with the P522R construct over the wild type. For comparison, the inflammation-related Ali5 D993G mutation exhibited a larger (2.27 ± 0.43 fold, mean value ± SD) increase in enzyme function upon EGF stimulation (Figure 3A).

**Figure 3.**
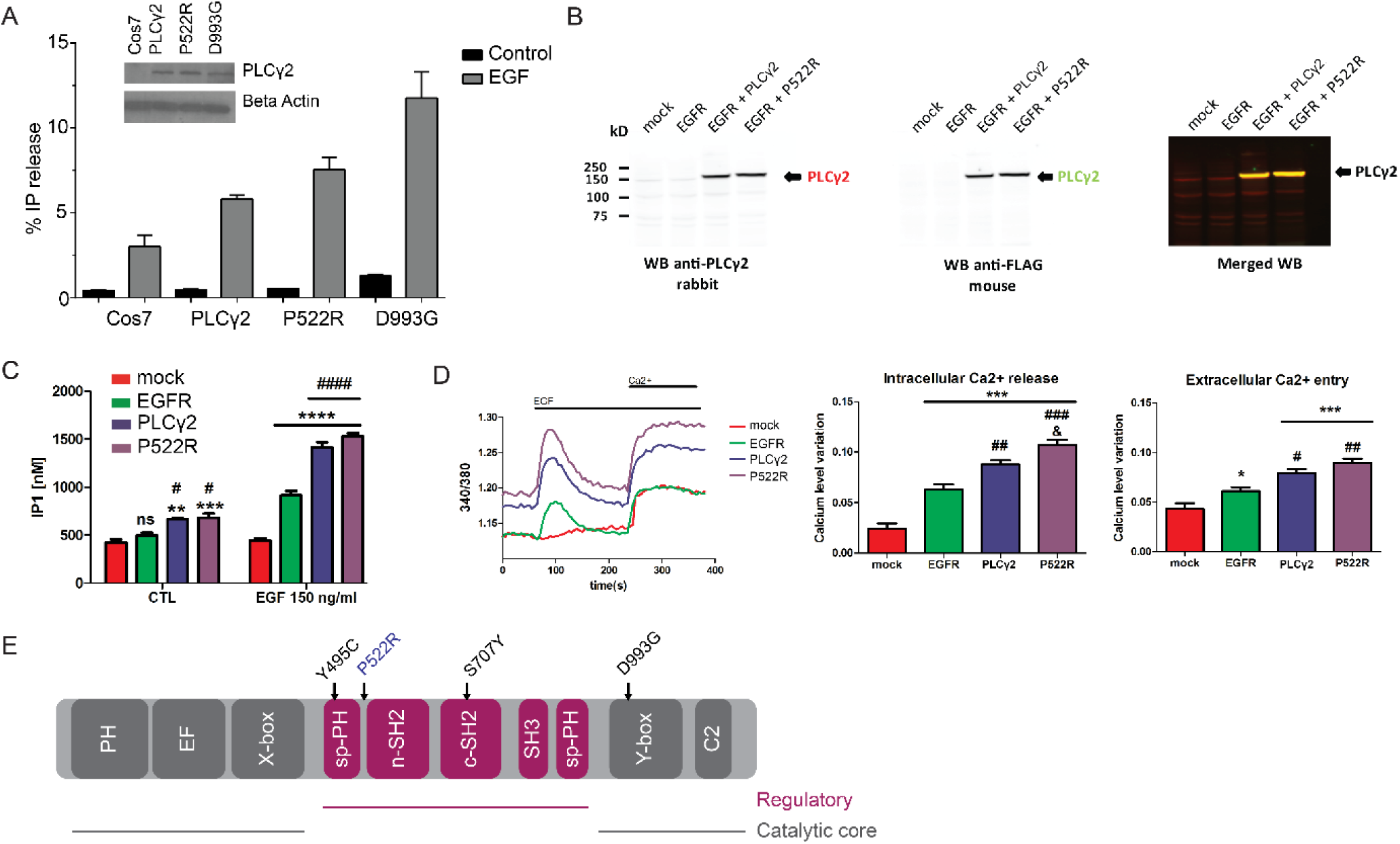
PLCγ2 protective variant P552R shows a slightly hypermorphic activity by increasing intracellular calcium release. A) PLC activity of the P522R PLCγ2 variant under basal and stimulated conditions. Measurement of % IP release in non-transfected COS7 cells (COS7), COS7 cells transfected with pTriEx4-PLCG2 (common variant; PLCG2 522P), construct with the AD associated, rare variant (522R) and construct with the D993G (Ali5) mutation. Western blotting was used to confirm equal expression (inset). SD is represented by error bars. The data are representative for 3 independent experiments. B) HEK293T were transfected with mock, EGFR, EGFR-PLCG2 or EGRF-PLCG2 P522R. Transfected cells where stimulated with EGF for quantification of IP1 by ELISA or for monitoring intracellular Ca2+ changes. C) Quantification of IP1 by ELISA IP-One. Transfected HEK293T cells were stimulated with EGF 150 ng/ml and analyzed for IP1 quantification. ELISA reading were averaged of ± standard error (n = 3, *:p < 0.05,**:p <0.01, ***:p <0.001, ****:p <0.0001, #:p <0.05, ### #:p <0.0001, Two-way ANOVA, Bonferroni multiple comparisons, *: compared to control, #: compared to EGFR). D) Transfected HEK293T cells loaded with Fura-2AM where stimulated with human EGF 150 ng/ml. Intracellular calcium level was recording by calculating the 340/380 nm excitation ratio at each 5 seconds. The net intracellular Ca2+ releases value was obtained by subtracting the average of 340/380 excitation ratio values between 0 and 50 seconds from the maximum 340/380 excitation ratio value between 60 and 200 seconds after the EGF stimulation. The net extracellular Ca2+ entry value was obtained by subtracting the average 340/380 excitation ratio values between 285 and 380 seconds from the average of 340/380 excitation ratio values between 215-235 sec after adding the extracellular calcium. Data in D were averaged of ± standard error (*:p < 0.05, **:p < 0.01,***:p < 0.001, #:p < 0.05, ##:p < 0.01, ###:p < 0.001, &:p < 0.05, One-Way ANOVA, Tukey’s Multiple Comparison Test, *: compared to mock, #: compared to EGFR, &: compared to PLCG2. The “n” is corresponding to an average of 45 reading from 3 independents transfection). E) Linear representation of the domains of PLCγ2 with locations of the point mutations (Y495C, Ali14; P522R, AD protective variant; S707Y, APLAID; D993G, Ali5)

IP3 is quickly degraded in IP1 which can be measured by ELISA. As a second functional endpoint, we therefore evaluated the enzyme activity of PLCγ2 P522R by measuring the cellular level of IP1 after activation of transfected epidermal growth factor receptor (EGFR) into HEK293T cells. In figure 3B, we confirmed expression of both the wild type and the P522R PLCγ2 variant. Stimulation of mock transfected cells with EGFR failed to increase IP1 level (Figure 3C). The expression of EGFR or EGFR + PLCγ2 significantly increased the amount of IP1 confirming an activation of the receptor and PLCγ2, PLCγ2 P522R enzyme activity was similar to PLCγ2, as measured by IP1 production (Figure 3C).

As a third endpoint of PLCγ2 function, we measured real time changes in intracellular Ca2+. No Ca2+ changes were seen in mock HEK293T cells stimulated with EGF (Figure 3D), confirming our previous ELISA IP-One results, and as previously observed (Zhou et al., 2012). Expression of EGFR induced significant intracellular Ca2+ changes after addition of EGF, as measured by increase of the 340/380 nm excitation ratio (Figure 3D, left panel). Quantification of the intracellular Ca2+ release showed a small but significant increase for PLCγ2 P522R variant as compared to wild type PLCγ2 (Figure 3D, middle panel). Depletion of the intracellular Ca2+ store activates a Ca2+ flux across the plasma membrane, referred to store-operated Ca2+ entry or capacitative Ca2+ entry (Putney, 1986; Putney et al., 2001). This Ca2+ flux was observed by exposing the cell to extracellular Ca2+ (Figure 3D, right panel). Ca2+ level monitoring show no significant differences in the extracellular Ca2+ entry between cells expressing EGFR + PLCγ2 and EGFR + PLCγ2 P522R (Figure 3D, right panel).

Taken together, these data are consistent with the PLCγ2 P522R variant conferring a small hypermorphic effect on enzyme function. The molecular mechanism by which this occurs is currently unknown. Other mutations in PLCG2 that lead to immune dysfunction are known to be activating (Koss et al., 2014). These mutations have been proposed to affect enzyme function by disruption of the autoinhibitory interface, or destabilisation of the regulatory domain that blocks enzyme inactivation. P522R is certainly located within a region of the enzyme that has a regulatory function (Figure 3D, Koss et al 2014), which is consistent with this polymorphism modulating enzyme activation.

In summary, we showed that PLCG2 is expressed in human and mouse brain immune cells and its expression is maintained in disease-associated microglia in the cerebral tissue of an APP mouse model. The rare coding Alzheimer’s protective variant shows a small hypermorphic activity upon stimulation in various orthogonal assays. Further experiments will need to address the functional consequences of the protective variant on immune cell phenotype. However, these findings allow us to speculate that weak lifelong activation of the PLCγ2 pathway might confer protection against developing AD, and provide evidence that a limited activation of this enzyme may have a beneficial therapeutic effect. These data, along with studies on TREM2 and other AD associated variants (Wang et al., 2015), appear to show consistent directionality that challenges long-standing dogma in the field that immune suppression is beneficial in the setting of AD. Indeed, emerging data indicates that variants which promote immune activation appear to be associated with decreased AD risk and conversely variants which functionally are associated with immune suppression are associated with increased risk.

## SUPPLEMENTARY FIGURES

**Supplementary Figure 1.**
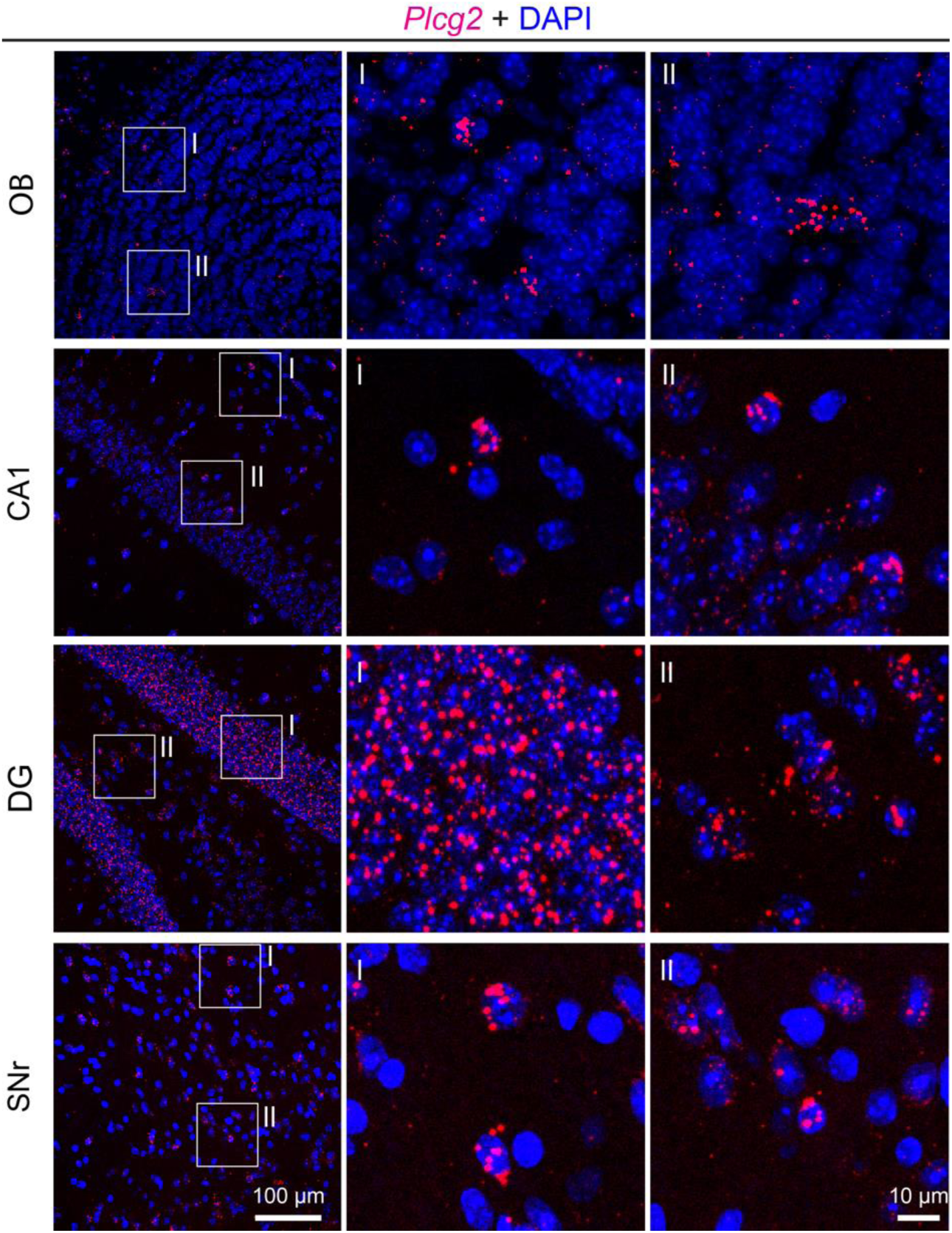
PLCγ2 antibodies and *in situ* hybridization on mouse brain tissue RNAScope for *Plcg2* in adult mouse brain (OB, olfactory bulb; CA1, hippocampal area CA1; DG, dentate gyrus; SNr, substantia nigra pars reticulata).

**Supplementary Figure 2.**
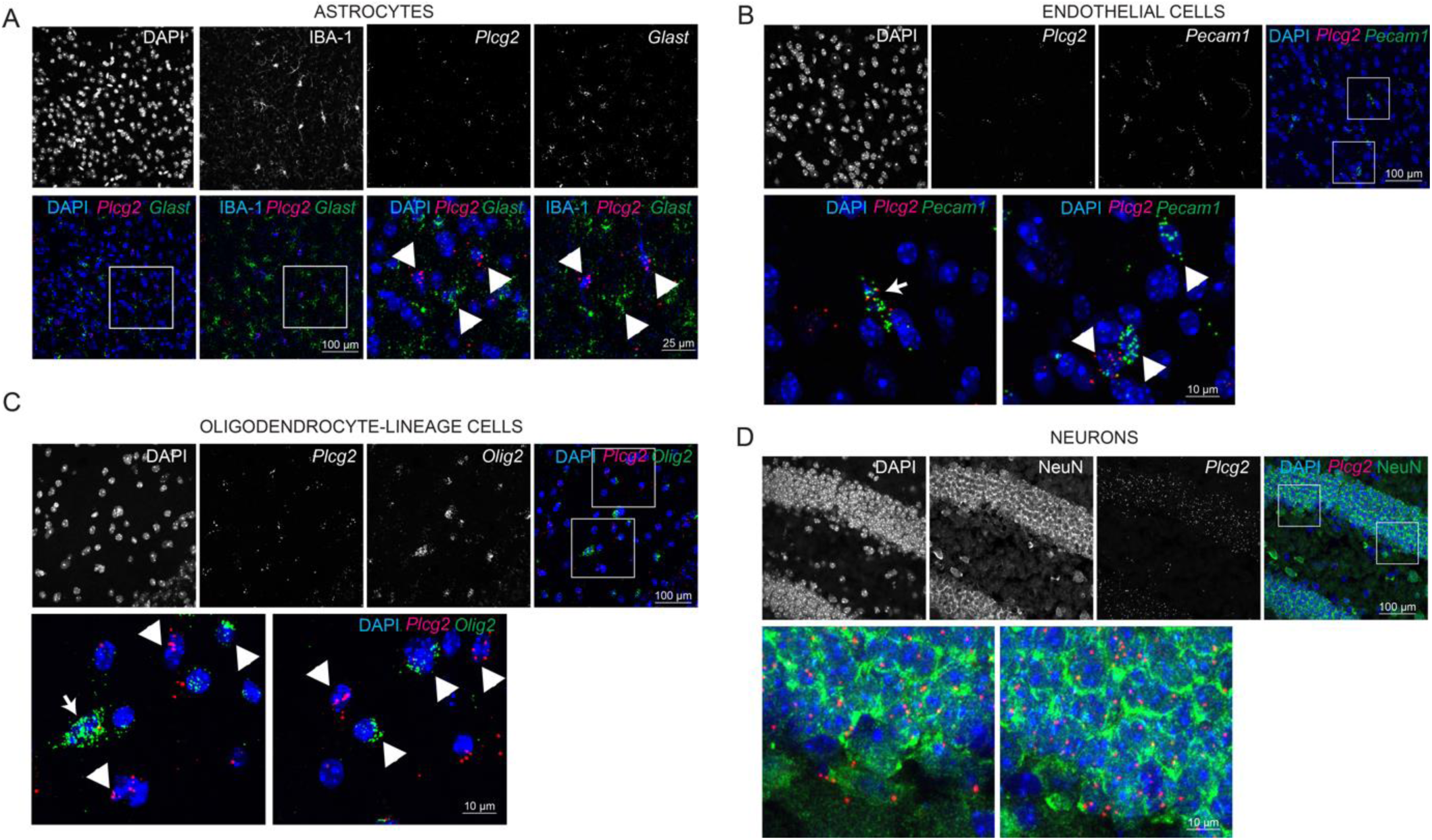
Characterisation of Plcg2 cell-type co-expression A) Multiplex RNAScope for *Plcg2* and *Glast* followed by IHC for IBA-1 on adult mouse cortex. Arrowheads indicate single-labelled cells. B) Multiplex RNAScope for *Plcg2* and *Pecam-1* on adult mouse cortex. Arrows point to co-expression of the two markers, arrowheads point to single-labelled cells. C) Multiplex RNAScope for *Plcg2* and *Olig2* on adult mouse cortex. Arrows point to co-expression of the two markers, arrowheads point to single-labelled cells. D) RNAScope for *Plcg2* followed by IHC for NeuN on dentate gyrus granule cell layer.

**Supplementary Figure 3.**
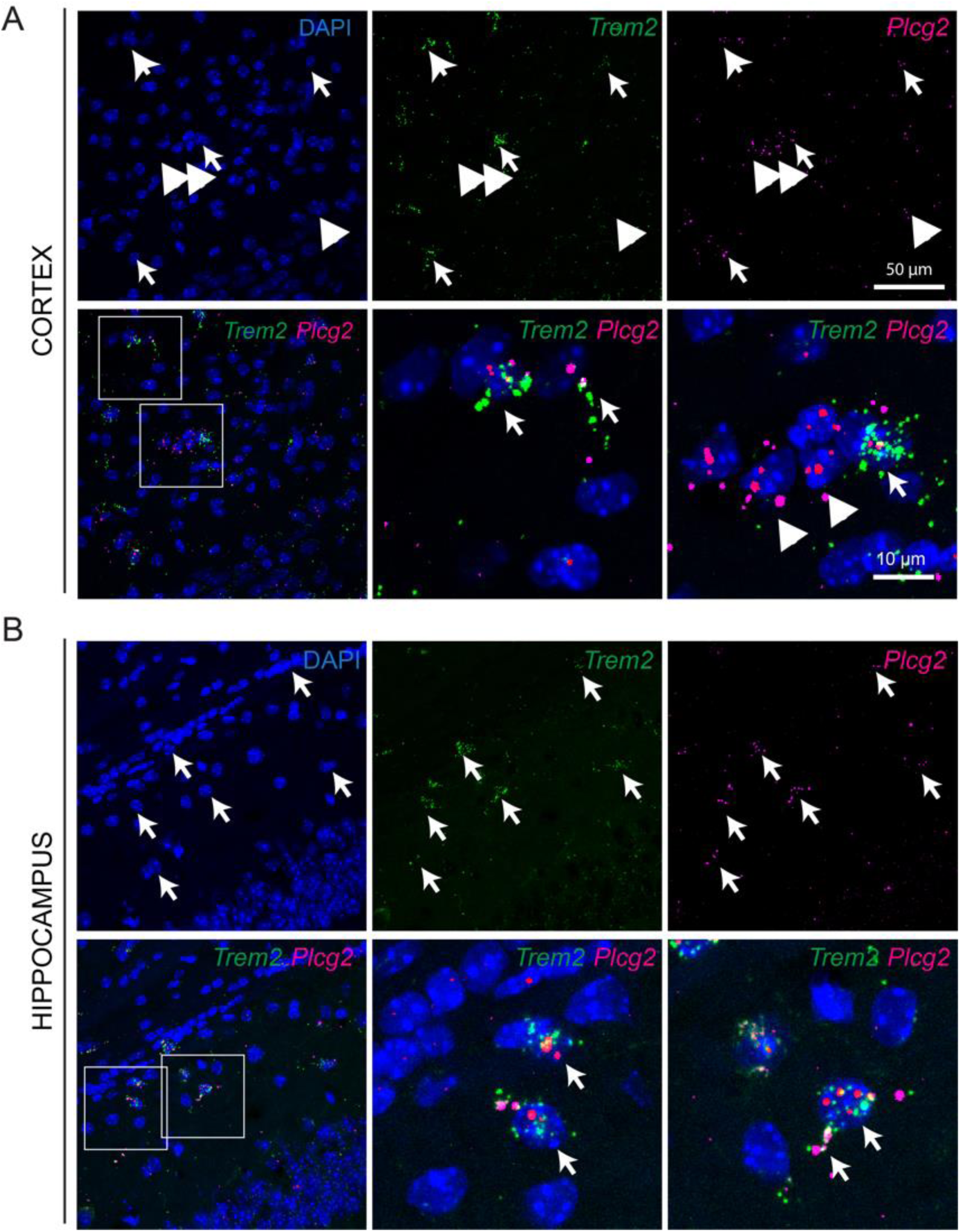
Plcg2 and Trem2 co-localisation analysis. A) Multiplex RNAScope for *Plcg2* and *Trem2* in the adult mouse cortex. Arrows point to co-expression, arrowheads indicate single-labelled cells. B) Multiplex RNAScope for *Plcg2* and *Trem2* in the adult mouse hippocampal CA1. Arrows point to co-expression, arrowheads indicate single-labelled cells. Note co-localisation in processes of putative microglia.

## MATERIALS AND METHODS

*Animals*. WT mice were maintained on a C57BL6 background at the Wolfson Institute for Biomedical Research in accordance with United Kingdom legislation (ASPA 1986).

TgCRND8 mice were maintained in-house by breeding APP transgenic males (carrying wild type RD gene) with C57B6/C3H F1 females (Envigo) (PMID: 11279122). These mice have florid AD-type Aβ plaque pathology in their forebrains, starting around 3 months of age. Animal procedures were approved by the University of Florida Institutional Animal Care and Use Committee. All animals were house grouped, under standard laboratory conditions (12:12 hours light/dark cycle, lights on at 0600 hours) with a room temperature of 21°C, and water and food available *ad libitum*.

*Tissue processing*, *immunohistochemistry (IHC)*, *and in situ hybridization (ISH)* were carried out as previously described (Magno et al., 2012, Rubin et al., 2010). Primary antibodies used were the following: rabbit anti-PLCγ2 (H160, Santa Cruz Biotechnologies sc-9015); rabbit anti-PLCγ2 (custom produced and purified by Pacific Immunology Corp (Ramona, CA) using the peptide sequence ‘INSLYDVSRMYV’); rabbit anti-Iba-1 (Wako).

Adult mice were perfusion-fixed with 4% paraformaldehyde (PFA), and the brain were dissected out and post-fixed overnight in 4% PFA. Samples were cryoprotected by overnight immersion in 20% sucrose, embedded in optimal cutting temperature (OCT) compound (Tissue Tek, Raymond Lamb Ltd Medical Supplies, Eastbourne, UK) and frozen on dry ice. 15 *μ*m cryosections were collected onto Superfrost slides and *in situ* hybridization (RNAScope) was carried out according to manufacturer’s instructions with the following catalogue probes: Mm-Plcg2 474781; Mm-Plcg1-C2 483531-C2; Mm-Olig2-C2 447091-C2; Mm-Pecam1-C3 316721-C3; Mm-Slc1a3-C2 430781-C2; Mm-Trem2-C2 404111-C2 (ACD, Biotechne). For some experiments *in situ* hybridization was followed by immunohistochemistry.

All sections were counterstained with Hoechst 33258 dye (Sigma, 1000-fold dilution) and the slides were mounted with Dako Fluorescence Mounting Medium (DAKO).

Confocal images (z stack height on average 10 μm, 1 *μ*m spacing) were taken on a Zeiss LSM 880 confocal microscope (Carl Zeiss AG) and processed for contrast and brightness enhancement with Photoshop (CS5, Adobe). A final composite was generated in Adobe Illustrator (CS5, Adobe).

### Human tissue processing

Post-mortem human frontal cortex formalin-fixed, paraffin-embedded, 5 μm thick sections were supplied by Queen Square Brain Bank for Neurological Disorders, UCL-Institute of Neurology (London, UK). The tissue is stored for research under a license from the Human Tissue Authority. Sections were deparaffinized in xylene, and rehydrated in graded ethanols (100%, 90% and 70%). Endogenous non-specific peroxidase activity was quenched by 0.03% H_2_O_2_ (Sigma-Aldrich) in methanol (1:100 dilution) for 5 minutes. Followed by 10 min antigen retrieval treatment. After blocking with 10% non-fat milk solution for 1 hour at room temperature, slides were incubated with primary antibodies overnight at 4°C (1:50). Sections were incubated with biotinylated anti-rabbit IgG (1:200, Sigma-Aldrich Ltd.) for 30 minutes at room temperature. After three 5 minutes washes in PBS sections were incubated with avidin-biotin complex horseradish peroxidase (ABC) reagent (Thermo Fisher Scientific) for 30 minutes. The reaction was developed with diaminobenzidine (DAB) activated by H_2_O_2_. Sections were counterstained with Mayer’s Haematoxylin (RAL Diagnostics). Sections were dehydrated though increasing concentrations of ethanol (70%, 90% and 100%) and cleared in xylene series. Finally, slides were mounted with DPX (Sigma-Aldrich) and cover slipped.

### Cell culture and transfections

Plasmids for the expression of full-length human pEGFPC1-PLCG2 constructs in mammalian cells have been described previously (Matsuda et al., 2001). PLCG2 is from Origene (ref. PMID: 25568344). EGFR construct is a gift from Axel Ullrich (Addgene plasmid # 65225). QuikChange PCR mutagenesis (Stratagene) was used to introduce the P522R point mutation in the PLCG2 WT variant. All mutants were fully sequenced to verify the fidelity of PCR.

COS7 cells were maintained at 37 ^°^C in a humidified atmosphere of 95% air and 5% CO2 in Dulbecco’s modified Eagle’s medium (DMEM) (Invitrogen) supplemented with 10% (v/v) fetal bovine serum (Invitrogen) and 2.5 mM glutamine. Prior to transfection cells were seeded into 6-well plates at a density of 2.5 x 10^5^ cells/well and grown for 16 h in 2 ml/well of the same medium. For transfection, 1.0 ug of PLCG2 plasmid DNA was mixed with 1 ul of PlusReagent^TM^ and 7 ul of Lipofectamine^TM^ (Invitrogen) and added to the cells in 0.8 ml of DMEM without serum. The cells were incubated for 3.5 h at 37 ^°^C, 5% CO2 before the transfection mixture was removed and replaced with DMEM-containing serum.

### Analysis of Inositol Phosphate Formation in Intact COS7 Cells

Inositol Phosphate formation was assessed as described in Everett et al., 2009.

Briefly, 24 h after transfection, the cells were washed twice with inositol-free DMEM without serum and incubated for 24 h in 1.5 ml of the same medium supplemented with 0.25% fatty acid free bovine serum albumin (Sigma) and 1.5 uCi/ml myo-[^2- 3^H]inositol (MP Biomedicals). After a further 24 h, the cells were incubated in 1.2 ml of inositol-free DMEM without serum containing 20 mM LiCl with or without stimulation with 100 ng/ml EGF (Calbiochem). The cells were lysed by addition of 1.2 ml of 4.5% perchloric acid. After incubating the samples on ice for 30 min, they were centrifuged for 20 min at 3700 x g. Supernatants and pellets were separated. The supernatants were neutralized by addition of 3 ml of 0.5 M potassium hydroxide/9 mM sodium tetraborate and centrifuged for a further 20 min at 3700 x g. Supernatants were loaded onto AG1-X8 200–400 columns (Bio-Rad) that had been converted to the formate form by addition of 2 M ammonium formate/0.1 M formic acid and equilibrated with water. The columns were washed three times with 5 ml of 60 mM ammonium formate/5 mM sodium tetraborate, and inositol phosphates were eluted with 5 ml of 1.2M ammonium formate/ 0.1 M formic acid. 5 ml Ultima-Flo scintillation fluid (PerkinElmer Life Sciences) was added to the eluates and the radioactivity quantified by liquid scintillation counting. The values represent total inositol phosphates. The pellets from the first centrifugation were resuspended in 100 ul of water and 375 ul of chloroform/methanol/HCl (200:100:15) was added. The samples were vortexed, and an additional 125 ul of chloroform and 125 ul of 0.1 M HCl were added.

After further vortexing, the samples were centrifuged at 700 x g for 10 min. 10 ul of the lower phase were placed in a scintillation vial with 3 ml of Ultima-Flo scintillation fluid and the radioactivity quantified by liquid scintillation counting. The obtained values correspond to radioactivity in inositol lipids. PLC activity is expressed as the total inositol phosphates formed relative to the amount of [^3^H]myo-inositol in the phospholipid pool. Because the differences in steady state labeling of inositol lipids are small (within 20%), this normalized PLC activity corresponds closely to PLC values expressed as total inositol phosphates.

### Western Blotting

PVDF membranes were blocking in TBS 0.5% casein a one hour at room temperature. Antibodies were diluted in TBS with 0.2% Tween-20 (TBS-T) and incubated one hour at room temperature. The membranes were washed with TBS-T and Western Blots analyzed with Odyssey infrared imaging system (LI-COR Inc., NE, USA).

### ELISA D-myo-inositol 1-phosphate

D-myo-inositol 1-phosphate (IP1) was quantified by ELISA IP-One (CISBIO US, Inc, USA) according to the manufacturer’s instructions. Briefly, transiently HEK293T co transfected with EGFR and PLCG2 were stimulated with human recombinant EGF (ThermoFisher, Carlsbad, USA) 150 ng/ml for 1 hour. Cell lysates were loaded in ELISA plate for IP1 quantification.

### Calcium flux assay

Intracellular calcium (Ca2+) changes were measured with FURA-2-AM (Invitrogen, USA) based on Zhu et al. (X. Zhu, M. Jiang, M. Peyton, G. Boulay, R. Hurst, E. Stefani, L. Birnbaumer, Cell 85 (1996) 661– 671.). Transiently transfected HEK293T cells attached to poly-L-lysine-treated plate were washed with HBSS (HBSS-Hepes buffer; 120 mM NaCl, 5.3mM KCl, 0.8 mM MgSO4, 1.8 mM CaCl2, 10 mM glucose,20 mM Hepes, pH 7.4) and loaded with Fura 2/AM(Sigma) for 20 minutes at room temperature in the dark. Cells were washed with HBSS and incubated with HBSS for 30 minutes at room temperature. The intracellular Ca2+ changes were measured by alternating the fluorescence excitation wavelengths 340 and 380 nm and emitted fluorescence at 510nm. Data acquisition was typically at 5-seconds intervals and data were presented as emitted fluorescence ratio 340/380.

## AUTHOR CONTRIBUTIONS

PW, LM, MK, JB, TEG and CBL designed the study. PM, LM, CBL, PC and MM performed the experiments. LM and PW wrote the manuscript. CBL, MK, TEG, PC and JB contributed to the manuscript. PW, TEG and PC acquired financial support.

### ACKNOWLEDGMENTS

We would like to thank our colleagues at the UCL ARUK-DDI for technical help and useful discussion; and Tammaryn Lashley, Queen Square UCL Brain Bank, for the provision of human tissue. This work was supported by Alzheimer’s Research UK; NIH grants U01 AG046139 R01 AG018454 and P50 AG047266, Florida Department of Health Grant 8AZ16

The authors declare no competing financial interests.

